# A specialized inhibitory function sharpens somatosensory hand representation and enhances the production and perception of fast multifinger movements in pianists

**DOI:** 10.1101/2024.01.23.576947

**Authors:** Masato Hirano, Yudai Kimoto, Sachiko Shiotani, Shinichi Furuya

## Abstract

Accurate control of fast, coordinated movements across multiple body parts characterizes experts’ skills, such as playing musical instruments. While performing such skillful movements, the somatosensory system is challenged to successively and in parallel process a large amount of somatosensory information originating from different body parts within a short period. Over decades, it has been posited that the cortical representations of distinct body parts are more isolated from each other in trained than untrained individuals. Several recent studies, however, have re-examined and failed to replicate it. Here, we provide compelling evidence that expert pianists possess a unique inhibitory function that isolates the somatosensory processing of different body parts in the somatosensory cortex (S1). A behavioural experiment demonstrated a superior ability to perceive fast multifinger movements in pianists than musically untrained individuals, suggesting the specialized neural process of somatosensory information originating from multiple fingers within a short period in pianists. A series of neurophysiological experiments demonstrated that pianists have a unique inhibitory function in the S1, which was activated by weak electrical stimulation to the ulnar nerve. This stimulation also increased the representational distance between fingers, which was assessed based on cortical activation patterns elicited by the passive finger movements. This indicates the strengthened independence of the individual finger representation in the somatosensory processes specifically in pianists. This stimulation also augmented both the perception and execution of the fast and complex multifinger sequential movements. In nonmusicians, neither the inhibitory effects on the somatosensory process nor enhancement of the perception of multifinger movements was induced by this stimulation. Together, these findings provide the first evidence of the experience-dependent plasticity of inhibition of the somatosensory system, which highlights its pivotal role in the isolated somatosensory processing of multiple body parts in trained individuals and enables them to control fast and complex multifinger movements.

## Introduction

Over centuries, the virtuosity of musicians and athletes has fascinated audiences. These dexterous motor skills commonly require fast and skilful control of multiple body parts. For example, pianists control bimanual finger movements with high temporal and force precision at high speeds to produce beautiful music(*1*). During these movements, afferent somatosensory signals from multiple fingers are input into the nervous system in a short period, which can interfere with each other. This is due to the architecture of the nervous system; first, cortical representation overlaps between multiple body parts in the sensorimotor cortex(*2*), and second, some sensory inputs from different body parts converge into identical neurons(*3*, *4*). How do experts solve this problem and perform fast, complex movements with high precision? Previous studies have demonstrated that the distance of cortical representation in the somatosensory cortex (S1) between fingers can be plastically adapted through training of the use of a robotic finger(*5*), is larger in string instrument players than in untrained individuals(*6*), and smaller in musicians who suffer from focal-hand dystonia(*7*). This suggests that the increased distance of cortical representation is one candidate mechanism to circumvent the interference of somatosensory information across different body parts. However, this idea is not fully validated since recent studies failed to replicate these findings; rather, they demonstrated unchanged cortical representation in the S1 in pianists(*8*) and dystonia patients compared with healthy participants(*9*).

Another possibility is the enhancement of inhibitory functions in the S1, which are known to play roles in isolating somatosensory information in both spatial and temporal dimensions. Specifically, the activity of inhibitory interneurons in the S1 contributes to sharpening the representation of each body part by inhibiting the representation of adjacent body parts(*10*) and enhancing temporal acuity by rapidly terminating excitatory activity(*11*). Therefore, inhibitory functions in the somatosensory processes would be of great significance in the control of fine motor skills in which multiple body parts are moved independently with fine temporal accuracy(*12*). These inhibitory functions can be studied by using paired-pulse (*13*) and dual somatosensory evoked potential (SEP) techniques(*14*, *15*). The former technique provides two somatosensory stimuli to the same nerve in succession, which suppresses the amplitude of the SEP evoked by the second stimulus depending on the interval of the two stimuli, called paired-pulse suppression (PPS). In contrast, the simultaneous application of two independent somatosensory stimuli suppresses the amplitude of the SEP compared to the arithmetic sum of the individual SEPs. The underlying mechanism is surround inhibition, in which somatosensory inputs from one body part inhibit the excitatory activities induced by inputs from neighbouring body parts(*14*, *16*). Previous studies have demonstrated that PPS is reduced by aging(*17*, *18*), and both PPS and surround inhibition are aberrant in patients with focal hand dystonia(*19*, *20*). However, to date, it remains unknown whether these inhibitory functions are enhanced through mastering dexterous motor skills and whether these functions are responsible for isolating cortical representations of multiple body parts.

Musicians are a unique population to address the issue, because playing musical instruments requires a high temporal accuracy of movements with highly coordinated and independent control of multiple digits(*1*, *21*). Here, we examined the experience-dependent plasticity of inhibitory activity in the S1 by comparing PPS and surround inhibition between expert pianists and musically untrained individuals. A series of neurophysiological and behavioural experiments including both univariate and multivariate analyses revealed that pianists possessed a unique somatosensory inhibitory function that sharpened the cortical somatosensory representation of their digits by the surround inhibition mechanism and that the activation of this inhibitory function improved both the perception and execution of fast and complex finger movements by expert pianists.

## Results

Four experiments were carried out to examine the experience-dependent plasticity of the inhibitory function of the S1 and to uncover the roles of the inhibitory function in the perception and production of complex finger movements. In total, 71 expert pianists and 57 musically untrained healthy individuals (nonmusicians) participated in the study. All pianists majored in piano performance at a musical conservatory and/or had extensive and continuous private piano training under the supervision of a professional pianist/piano professor. Nonmusicians had no experience practising musical instruments for over 5 years. All experimental procedures were carried out in accordance with the Declaration of Helsinki and were approved by the ethics committee of Sony Corporation.

### Experiment 1

We first conducted a behavioural experiment to confirm that skilled pianists can perceive the fast multifinger movements more accurately than nonmusicians. Fifteen expert pianists (23.1±3.6 years old [mean±SD], 11 females) and 15 nonmusicians (24.2±3.1 years old [mean±SD], 9 females) participated in this experiment and were asked to perform a perceptual test. Participants wore an exoskeleton robot hand(*22*) on their right hand during this test. The robot hand can produce flexion and extension movements on the metacarpophalangeal joint of five digits. In this test, second to fifth digits were flexed passively with a randomized order except for the order of 2-3-4-5 and 5-4-3-2 (i.e., one of the 22 possible sequences was randomly chosen, 2: index finger, 3: middle finger,4: ring finger, 5: little finger). The participants were asked to identify the order of passive finger movements and to respond to the perceived order by typing on a keyboard with their left hand using the same fingering as the perceived one. The interval between onsets of each finger flexion in the sequential passive movement varied across trials, which was determined by an adaptive Bayesian staircase method (ZEST)(*23*) to quantify the perceptual threshold at which a participant was able to identify the order of a sequence of passive finger movements with 80% accuracy. Figure 1 shows the perceptual threshold obtained from the pianists and nonmusicians. We found a significant difference in the threshold between the two groups (t=-5.09, p<0.01), indicating that the pianists have superior ability to perceive the order of fast multifinger movements compared with nonmusicians.

**Figure 1.**
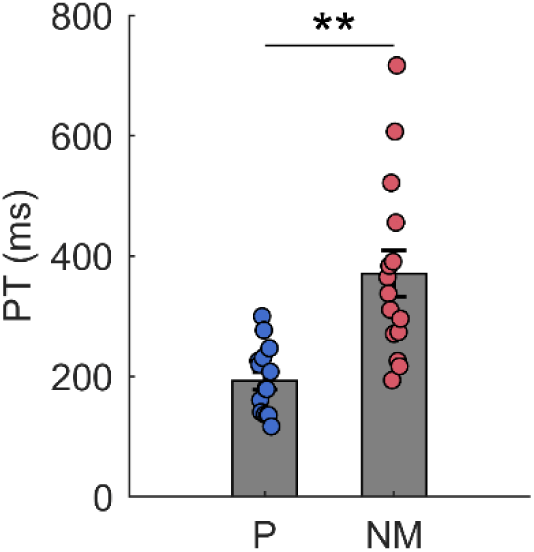
Perceptual threshold for identifying the order of passive multifinger movements obtained from pianists (P) and nonmusicians (NM). The vertical axis represents the timing interval between successive finger flexion motions at which a participant was able to identify the order of passive movements with 80% accuracy. Coloured dots indicate data of individual participants. Mean±SEM. **: P vs. NM, p<0.01.

### Experiment 2

The purpose of Experiment 2 was to examine the difference in somatosensory inhibitory function between pianists and nonmusicians. In total, 14 expert pianists (22.5±3.4 years old [mean±SD], 12 females) and 14 nonmusicians (22.3±1.7 years old [mean±SD], 11 females) participated in this experiment.

### Weak electrical stimulus on the ulnar nerve modulates subsequent somatosensory processing differently between pianists and nonmusicians

The inhibitory function in the S1 was quantitatively assessed according to the PPS of SEPs recorded from the left somatosensory area (electrode CP3, reference electrode: Fz) using an electroencephalogram (EEG) system. The right ulnar nerve and median nerve at the wrist were stimulated by applying an electrical pulse (square wave, duration 1 ms) using a constant current stimulator to evoke the SEP. Stimuli were applied either as a single pulse or as paired pulses with interstimulus intervals (ISIs) of 5, 10, 20, 30, 40, and 50 ms to examine the effects of a preceding somatosensory stimulus on the subsequently evoked SEP. We set the intensity of the first stimulus, termed the conditioning stimulus (CS), at just below the perceptual detection threshold (PT), which is determined as the lowest electrical current at which the subject reported sensation (i.e., CS-PT), and at just below the motor threshold (i.e., CS-MT). The second stimulus, termed the test stimulus (TS), was set at an intensity just below the motor threshold to evoke a clear SEP. Figure 2A illustrates typical waveforms of the ulnar and median nerve SEPs obtained from a pianist. In most participants, the SEP waveform was characterized by clear responses at latencies of approximately 20 ms (N20) and 25 ms (P25). One pianist and one nonmusician had an N20-P25 peak-to-peak amplitude less than 0.5 μV, so we excluded these two participants from further analyses. There were no significant differences in the N20-P25 peak-to-peak amplitude between pianists and nonmusicians in either nerve when the SEP was evoked by the single TS (median: pianist=4.80±1.05 μV, nonmusician: 3.60±0.59 μV, t=-0.98, p=0.34; ulnar: pianist=4.16±0.63 μV, nonmusician: 3.44±0.55 μV, t=1.00, p=0.33).

**Figure 2.**
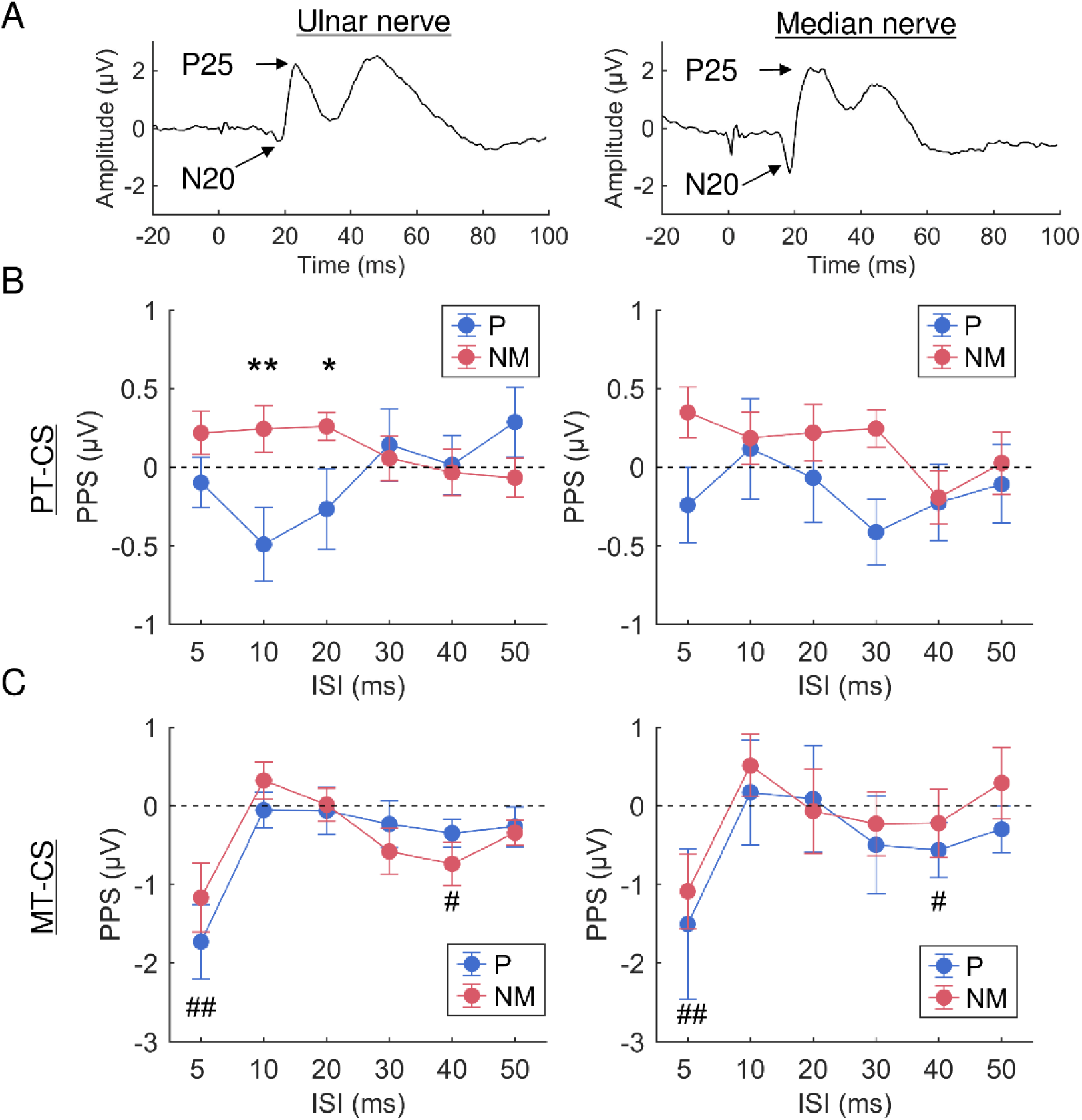
Effects of a leading somatosensory stimulus on the subsequent somatosensory processes. A: Typical SEP waveforms evoked by electrical somatosensory nerve stimulation. B, C: Effect of a conditioning stimulus on the N20-P25 amplitude of SEP. The horizontal axis indicates the ISI between the conditioning and test stimuli. The vertical axis represents the difference in the N20-P25 amplitude between the test stimulus only and paired-pulse conditions, an index of the PPS. The intensity of the conditioning stimulus was set just below the PT (B) and just below the MT (C). Blue and red lines represent results obtained from pianists and nonmusicians, respectively. The left and right panels represent the results obtained when the ulnar and median nerves were stimulated, respectively. Mean±SEM. *, **: pianist vs. nonmusician, p<0.05, p<0.01; ##, #: one-sample t test (vs. 0), p<0.01, p<0.05.

Figure 2B illustrates the PPS across different ISIs for ulnar and median nerve stimulation when the intensity of the CS was set just below the PT. The PPS was measured as the difference in the N20-P25 peak-to-peak amplitude between the paired-pulse SEP and the single-pulse SEP, which was evoked by only applying the TS. We first modelled the data using a generalized linear mixed-effects (GLME, Gaussian distribution, identity link function) model with the group, ISI, nerve, and their interactions as fixed effects; the participant, participant*ISI interaction, and participant*nerve interaction as random intercept effects; and the N20-P25 amplitude of a single-pulse SEP as the covariate. Type II Wald chi-square tests on the GLME results revealed a significant effect of the second-order interaction (group*ISI*nerve: χ^2^=11.23, p<0.05), indicating that the effects of a weak preceding stimulation on subsequent somatosensory processes varied depending on the group, ISI, and nerve. To simplify the results, we reanalysed the data obtained from the median nerve and ulnar nerve stimulation conditions separately. For the median nerve stimulation condition, PT-CS did not modulate the N20-P25 amplitude for all ISIs in either group (group: χ^2^=0.97, p=0.33; ISI: χ^2^=5.32, p=0.38; interaction: χ^2^=5.83, p=0.32). In contrast, for the ulnar nerve stimulation condition, PT-CS modulated the N20-P25 amplitude differently between the pianists and nonmusicians. The GLME model yielded a significant interaction between the group and ISI factors (χ^2^=14.57, p=0.01) on the PPS. The post hoc comparison using the estimated marginal means with Holm’s adjustment revealed a significant difference in the PPS between the two groups at ISIs of 10 ms (t=-2.88, p<0.01) and 20 ms (t=-2.01, p=0.04). Specifically, the SEP was inhibited and facilitated by PT-CS in the pianists and nonmusicians, respectively.

Figure 2C illustrates the effects of MT-CS on the N20-P25 amplitude evoked by the TS across different ISIs for ulnar and median nerve stimulation. A GLME model (fixed effects: group, ISI, nerve, and their interactions; random effects: participant, participant*ISI interaction, and participant*nerve interaction, covariate: amplitude of single-pulse SEP) yielded a significant fixed effect of the ISI (χ^2^=53.49, p<0.01) but not the group (χ^2^=0.37, p=0.54), nerve (χ^2^=0.1.94, p=0.16), or their interaction effects (all, p>0.05). Independent-sample t tests on the PPS revealed significant negative values at ISIs of 5 ms (t=-4.48, p<0.01) and 40 ms (t=-2.92, p<0.05). This indicates that a clear PPS was observed when the CS intensity was set above the MT, while the PPS did not differ between the two groups or between the nerves.

Taken together, these results demonstrated a specialized inhibitory function in the somatosensory processes in pianists that is activated specifically by weak ulnar nerve stimulation.

### Experiment 3

Experiment 2 demonstrated that weak somatosensory stimulation of the ulnar nerve differently modulates the subsequent somatosensory processes between pianists and nonmusicians. The purpose of Experiment 3 was to examine the functional roles of this effect in the perception of complex movements involving multiple fingers. In total, 14 expert pianists (22.6±1.8 years old [mean±SD], 11 females) and 14 nonmusicians (22.4±1.9 years old [mean±SD], 11 females) participated in this experiment.

### Replication of the different effects of weak ulnar nerve stimuli on subsequent somatosensory processing between pianists and nonmusicians

First, we examined the effects of weak ulnar nerve stimulation on subsequent somatosensory processing using EEG, as in Experiment 2, to test whether the results observed in the former experiment could be replicated in other participants. We recorded the SEP using the paired-pulse paradigm with ISIs of 5, 10, 20, 30, 40, and 50 ms. The CS intensity was set just below the PT, and the TS intensity was set just below the motor threshold. Figure 3A shows the PPS at different ISIs in the two groups. A GLME model (Gaussian distribution, link function: identity) yielded a significant fixed effect of group but not ISI or their interaction on the PPS (group: χ^2^=6.41, p=0.01; ISI: χ^2^=3.53, p=0.62; interaction: χ^2^=4.68, p=0.47). Consistent with the results of Experiment 1, the pianists generally exhibited an inhibitory effect, whereas nonmusicians exhibited a facilitatory effect. The strongest inhibitory effect in pianists was observed if the ISI was 10 ms.

**Figure 3.**
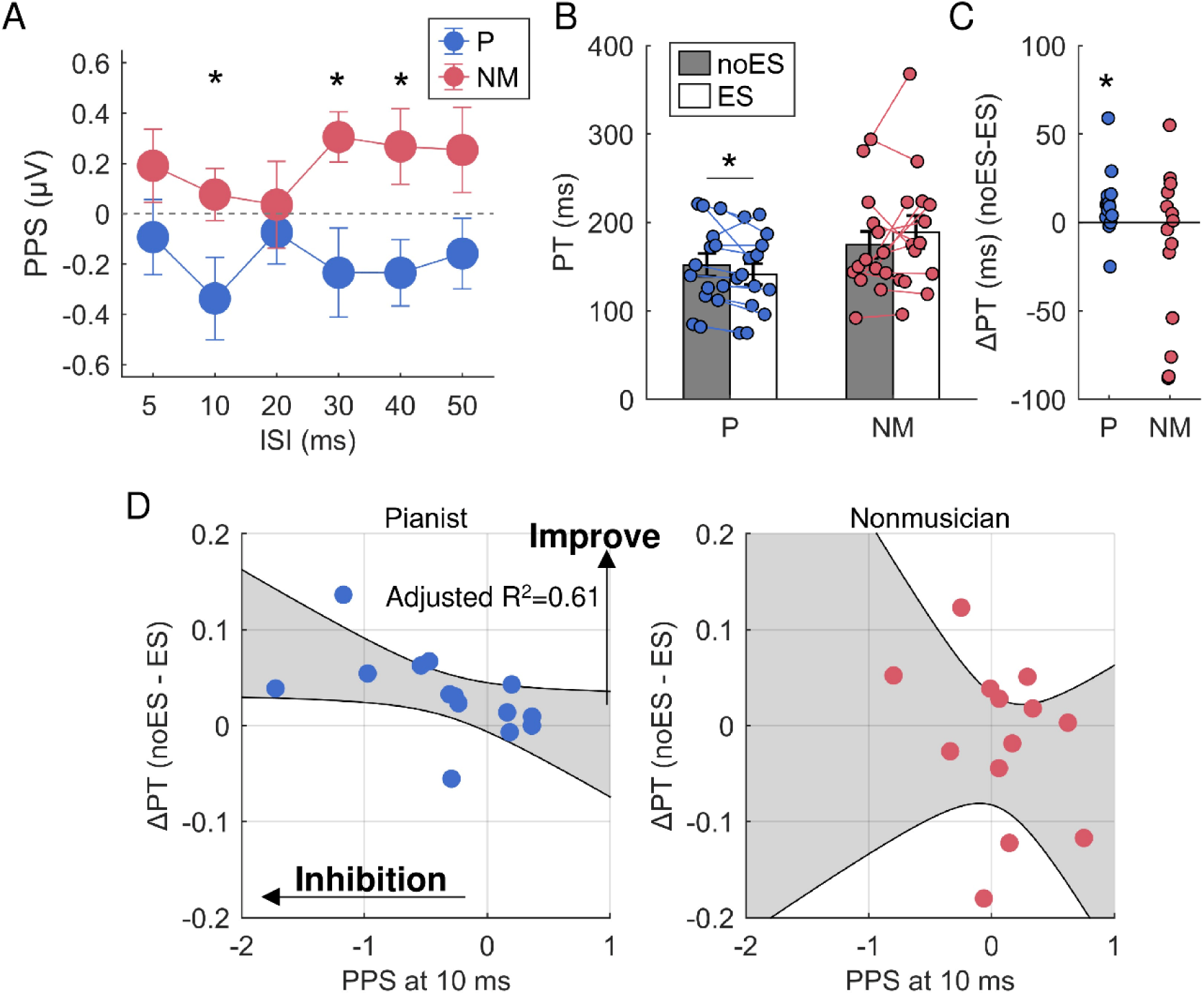
Effects of weak ulnar nerve stimulation on somatosensory perception. A: PPS assessed by paired-pulse SEP with weak ulnar nerve stimulation. *: pianist vs. nonmusician, p<0.05. B: Effect of weak ulnar nerve stimulation on somatosensory discrimination perception. The vertical axis represents the timing interval between each finger flexion at which a participant was able to discriminate the order of a sequence of passive finger movements with 80% accuracy. The left and right show the data obtained from pianists (P) and nonmusicians (NM). Coloured thin lines indicate individual data. The filled box represents the discrimination threshold when no electrical stimulation was provided, whereas the white box represents the threshold when weak ulnar nerve stimulation was provided in the test. Mean±SEM. *: with ES vs. without ES, p<0.05. C: The difference in the discrimination threshold between the ES and noES conditions. Each dot represents individual data. D: The correlation between the change in the discrimination threshold by the provision of ulnar nerve stimulation and the PPS at the ISI of 10 ms in pianists (left) and nonmusicians (right). The shaded area represents the 95% confidence interval. Note that the ΔPT was log-transformed.

### Weak somatosensory stimuli improve the perception of complex finger movements in pianists

In the former neurophysiological experiments, we found that the weak ulnar nerve stimulation activated the inhibitory function within the somatosensory processes. To examine the functional roles of this inhibitory activity on the perception of fast and complex finger movements, we performed a somatosensory perception test while the participants received the weak ulnar nerve stimulation. The participants wore the exoskeleton robot hand on their right hand during the perception test. In this test, the finger joints of the robot hand were flexed passively with one of two predetermined sequences (1-3-2-5 or 1-3-5-2; 1: thumb, 2: index finger, 3: middle finger, 5: little finger). We used the different procedure from the perceptual test used in Experiment 1 to minimize possible confounding effects of the left-hand dexterity and/or working memory on the performance. The participants were asked to identify in which sequence their fingers were moved by the robot hand (i.e., two-alternative forced choice task). The interval between onsets of each finger flexion in the sequential passive movement varied across trials, which was determined by the ZEST. The participants performed the perception test under two different conditions. The first condition was a weak electrical pulse (intensity just below the PT, 1 ms square pulse) applied to the ulnar nerve at the onset of each passive finger flexion (i.e., ES condition; the stimulation was applied 4 times in a single passive sequential movement because the sequence used in this test consisted of 4 successive finger flexions). In the second condition, electrical stimulation was not provided (i.e., noES condition). Figures 3B and 3C show the group means of the discrimination thresholds for the two conditions in the two groups and the difference in the threshold between the two conditions (i.e., ΔPT) in both groups, respectively. A GLME model (gamma distribution, link function: log; fixed effects: group, condition, and their interaction; random effects: participants) yielded a significant interactive effect between the group and condition on the discrimination threshold (χ^2^=7.61, p<0.01). In most pianists, the ΔPT was greater than zero (Fig. 3C), indicating that the PT in the ES condition was lower than that in the noES condition. A post hoc test revealed that the perceptual threshold was significantly reduced by the stimulation (z ratio=-2.03, p=0.04) in pianists but not in nonmusicians (z ratio=1.86, p=0.06). We also found that the PPS at an ISI of 10 ms was significantly correlated with the difference in the discrimination threshold between the two conditions in pianists (Fig. 3D left, robust regression coefficient: r=-0.84, adjusted R^2^=0.61, p<0.01) but not in nonmusicians (Fig. 3D right, r=-0.22, adjusted R^2^=-0.04, p=0.48). This suggests that the inhibitory activity evoked by weak ulnar nerve stimulation enhances the perception of multifinger movements specifically in pianists.

### Experiment 4

Experiments 2 and 3 demonstrated that weak ulnar nerve stimulation activates unique somatosensory inhibitory activity and enhances the perception of a fast and complex multifinger movement specifically in pianists. However, how the activation of the inhibitory function enhances perception remains unclear. Here, we examined the hypothesis that the weak ulnar nerve stimulation used in the present study activates neural surround inhibition (SI) circuits connected to the ulnar nerve, which spatially sharpens the representation of digits in the somatosensory processes. Fourteen pianists (22.0±2.2 years old [mean±SD], 10 females) and 14 nonmusicians (23.8±3.4 years old [mean±SD], 8 females) participated in this experiment.

### Weak ulnar nerve stimulation sharpens finger representation in the sensorimotor area in pianists

Previous studies using fMRI demonstrated that passive finger movements elicit unique spatial activity patterns in the S1 for each digit(*24*, *25*). On the basis of this observation, in this experiment, we quantified the dissimilarity in the scalp distribution patterns of the SEP amplitude evoked by passive finger movements between digits and examined the effect of weak ulnar nerve stimulation on it. Participants wore the exoskeleton robot hand on their right hand, and the robot hand had the metacarpophalangeal joint of one of four fingers flexed passively and had the thumb at the carpometacarpal joint rotated passively from 0 to 45 degrees at a constant speed of 450 degrees/s. Five hundred milliseconds after the onset of each passive movement, the robot moved these joints from 45 to 0 degrees at the same speed used for flexion. In another condition, the ring and little fingers were simultaneously flexed. The order of passive flexion across digits was randomized, and the interflexion interval was at least 2 seconds. The number of passive movements was 200 for each digit. In half of the trials, a weak electrical pulse (intensity just below the PT, 1 ms square pulse) was applied to the ulnar nerve at the onset of each passive flexion (i.e., ES condition), whereas no electrical stimulation was provided in the remaining trials (i.e., noES condition). The EEG was recorded from the whole brain using a 64-sensor system throughout the experiment.

To obtain SEPs, preprocessed EEG data were first segmented into epochs that ranged from −200 ms to 400 ms relative to the onset of passive flexion of each digit and then computed the SEPs for each digit and condition using the regression-based analysis. Then, we calculated the cross-validated Euclidean distance of prewhitened spatial patterns (i.e., across channels) of the SEP amplitude at each sample of the epoch between all possible pairs of digits (detailed methods are described in the Methods section) as a dissimilarity index. This yielded a single time course of dissimilarity for each digit pair. The resultant dissimilarity time courses were then averaged to obtain a single time course that represented the average interdigit dissimilarity across all five digits. SEPs obtained from all sensors (i.e., 64 sensors) were used for this analysis. Figure 4A shows the group mean time course of the interdigit dissimilarity of SEP spatial patterns for each condition in pianists and nonmusicians. Generally, the interdigit dissimilarity increased approximately 60 ms after the onset of passive flexion. Interestingly, weak ulnar nerve stimulation increased the interfinger dissimilarity in the pianists but not in the nonmusicians. The cluster permutation test for interactive effects between the group and condition factors revealed that there was a cluster that showed a significant interactive effect that extended from 108 ms to 132 ms after the onset of passive flexion (cluster t value = 20.09, p=0.02). We found that the interdigit dissimilarity was higher in the ES condition than in the noES condition in pianists but not in nonmusicians. Figure 4B shows the representational dissimilarity matrix consisting of the dissimilarity values of all possible pairs of digits and conditions observed at 108-132 ms after the onset of passive flexion. To visualize this, we applied classical multidimensional scaling to the group-averaged representational dissimilarity matrix. Figure 4C shows the two-dimensional map of the relationship of dissimilarity of spatial SEP amplitude patterns across digits. This map shows the enlarged distance of somatosensory representation between digits when applying weak ulnar nerve stimulation in pianists. This indicates that the inhibitory activity evoked by the weak ulnar nerve stimulation sharpens the individual finger representation in the early somatosensory process specifically for pianists.

**Figure 4.**
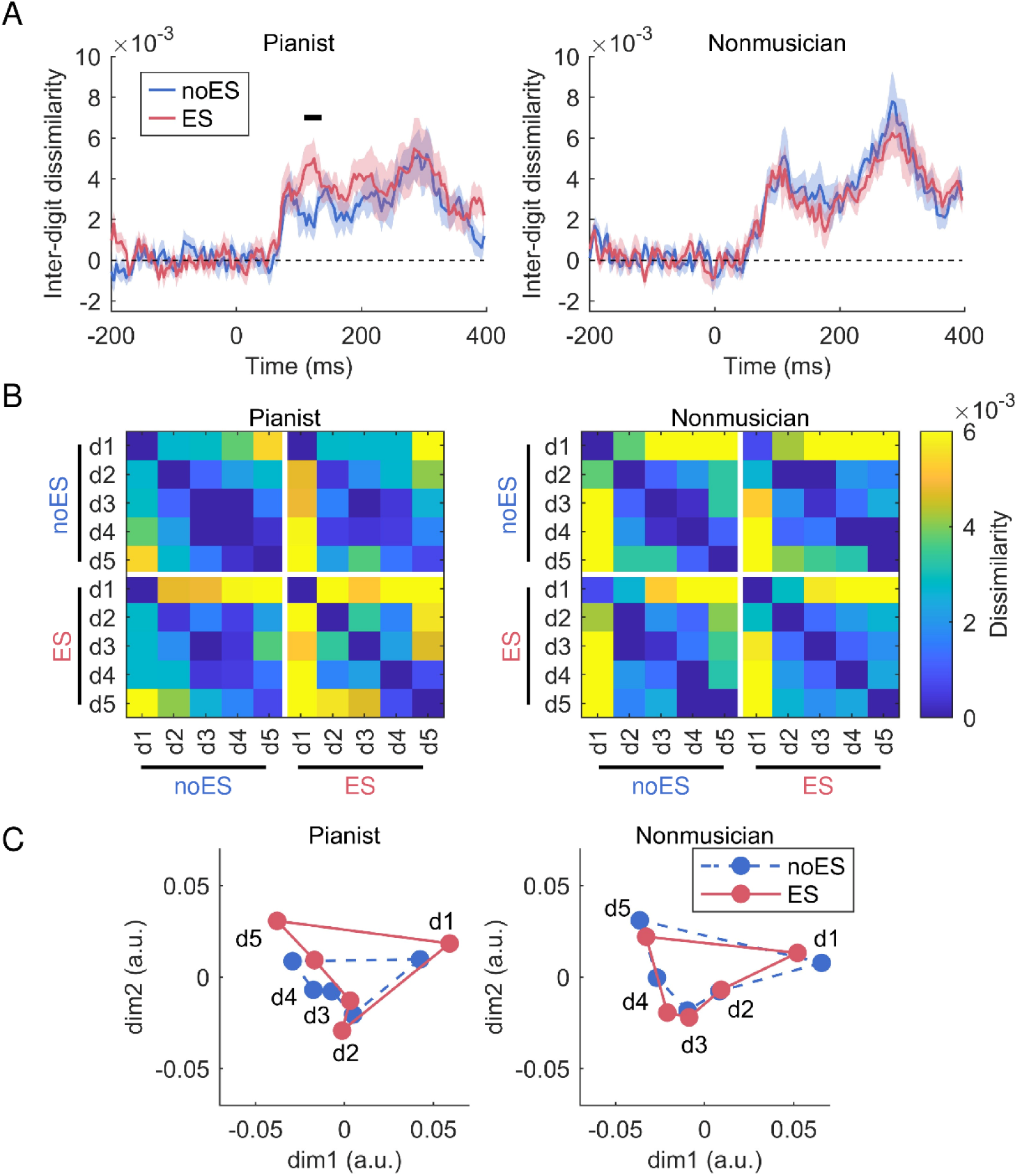
Effects of weak ulnar nerve stimulation on the subsequent proprioceptive processes. A: Effects of weak ulnar nerve stimulation on the average distance of spatial SEP amplitude patterns evoked by passive flexion of the MCP joint between two digits. The vertical axis represents the dissimilarity of spatial SEP amplitude patterns between two digits averaged across all possible pairs of digits. The blue lines represent the dissimilarity values obtained from the condition in which no electrical stimulation was provided, whereas the red line displays those obtained from the condition in which weak ulnar nerve stimulation was provided at the onset of the passive flexion of each digit. The horizontal line in the left panel indicates a cluster that shows significant differences in the dissimilarity between the two conditions (p<0.05). The shaded area represents the SEM across participants. B: Averaged representational dissimilarity matrices (RDMs) across participants and the time within the significant cluster. The left and right panels show the RDMs obtained from pianists and nonmusicians, respectively. Each cell represents the dissimilarity of spatial SEP amplitude patterns between each corresponding pair of digits. C: 2D depiction of the data in (B), using multidimensional scaling (MDS). MDS projects the higher-dimensional RDM into a lower-dimensional space while preserving the interdigit dissimilarity as accurately as possible. Blue dotted and red solid lines represent the data obtained from noES and ES conditions, respectively. Note: MDS plots are purely for visualization purposes and were not used for statistical analysis.

### Weak ulnar nerve stimulation enhances surround inhibition in the S1

To examine whether weak ulnar nerve stimulation activates the SI circuits in the S1, we compared the SEP amplitude evoked by single-finger movement and the simultaneous movement of two fingers. Figure 5A shows the SEP waveforms obtained from the CP3 electrode evoked by the passive flexion of the ring and little fingers from one representative pianist and nonmusician. The right panels show the SEPs when weak ulnar nerve stimulation was administered at the onset of passive movements, and the left panels show the SEPs without stimulation. The yellow lines represent an arithmetic sum of the two individual SEPs (i.e., sum SEP), and the purple lines indicate the SEP evoked by the simultaneous passive flexion of the ring and little fingers (i.e., dual SEP). Figure 5B shows the amplitude of the first component (i.e., P1 component) of the sum SEP, dual SEP, and the difference between the sum and dual SEPs in both groups. We found a significant interactive effect between the group and condition factors on the difference in the P1 amplitude between the sum and dual SEPs (GLME, Gaussian distribution, identity link function, fixed effects: group, condition, and their interaction; random effects: participant; group*condition: χ^2^=4.34, p=0.04). A post hoc test revealed that the difference in the P1 amplitude between the sum and dual SEPs was larger in the stimulation condition than in the no-stimulation condition in pianists (t=-2.14, p=0.04) but not in nonmusicians (t=0.81, p=0.43). This supports our hypothesis that weak ulnar nerve stimulation activates the surround inhibition circuits of the S1 in pianists.

**Figure 5.**
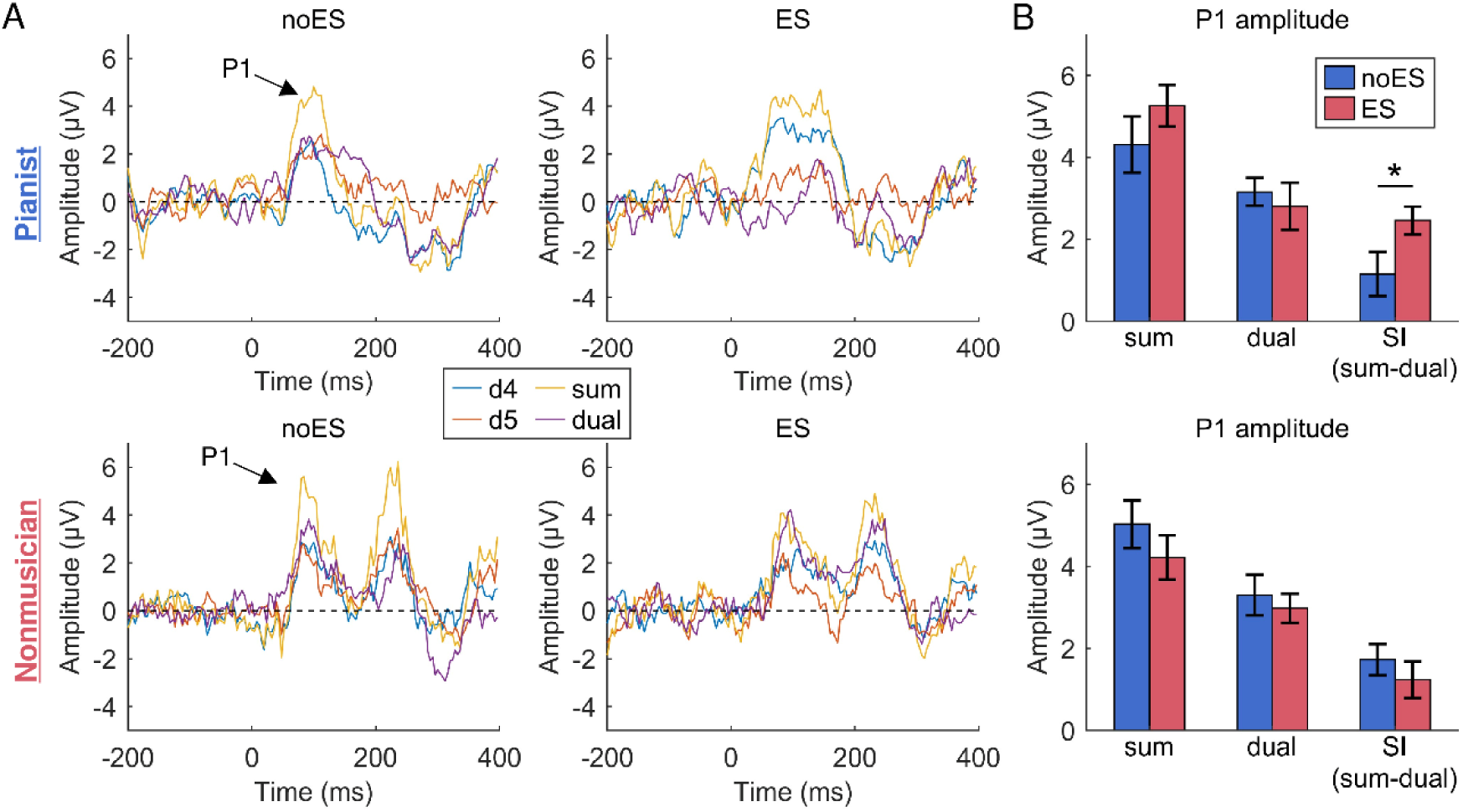
Effects of weak ulnar nerve stimulation on the surround inhibition (SI) of proprioceptive processes. A: Typical EEG waveforms evoked by passive finger flexion in a representative participant in each group. The blue and red lines represent the SEP evoked by the ring (d4) and little (d5) fingers, respectively. The yellow line indicates the arithmetic sum of the two SEPs (sum), and the purple line displays the SEP evoked by simultaneous flexion of d4 and d5 (dual). The upper and lower panels show the results obtained from a pianist and a nonmusician, respectively. We quantified the SI as the difference in the amplitude between the sum and dual SEPs. B: The group average of the amplitude and SI of the P1 component of sum and dual SEPs in both groups and in both conditions. Mean±SEM. * noES vs. ES, p<0.05.

### Experiment 5

The final experiment examined whether the inhibitory activity evoked by the weak ulnar nerve stimulation enhances the production of fast and complex sequential finger movements performed by pianists. Fourteen pianists (23.0±3.3 years old [mean±SD], 11 females) participated in this experiment. The pianists were asked to strike piano keys using their right hand in the following order: 1-3-4-2-3-5 (1: thumb, 2: index finger, 3: middle finger, 4: ring finger, 5: little finger). They repeated the sequence for 20 seconds as fast and accurately as possible during a trial. During this task, electrical pulses (1 ms square wave) with an intensity just below the PT were delivered to the right or left ulnar nerve at a frequency of 10 Hz. For the control condition, no stimulation was provided during the task.

Figure 6A shows the number of erroneous strikes due to incorrect fingering (i.e., mistouches) made during the task in each condition. A GLME model with a Poisson distribution (link function = log; fixed effects: condition; random effects: participants, participants*condition) identified a significant main effect of condition on these data (χ^2^=13.16, p<0.01). A post hoc test revealed that the number of mistouches was significantly smaller when electrical stimulation was applied to the right ulnar nerve than in the other two conditions (right vs. left: z-ratio=-2.64, p=0.02; right vs. no stimulation: z-ratio: −3.52, p<0.01; left vs. no stimulation: z-ratio=-0.89, p=0.38). This reduction in the number of mistouches was not due to a reduction in keystroke speed because the interkeystroke interval (IKI) for correctly produced sequences did not differ across conditions (Figure 6B, GLME, gamma distribution, log link function, fixed effects of the condition factor: χ^2^=0.07, p=0.97). Thus, this experiment demonstrated that weak somatosensory stimulation improved the fast and accurate execution of complex sequential finger movements in expert pianists.

**Figure 6.**
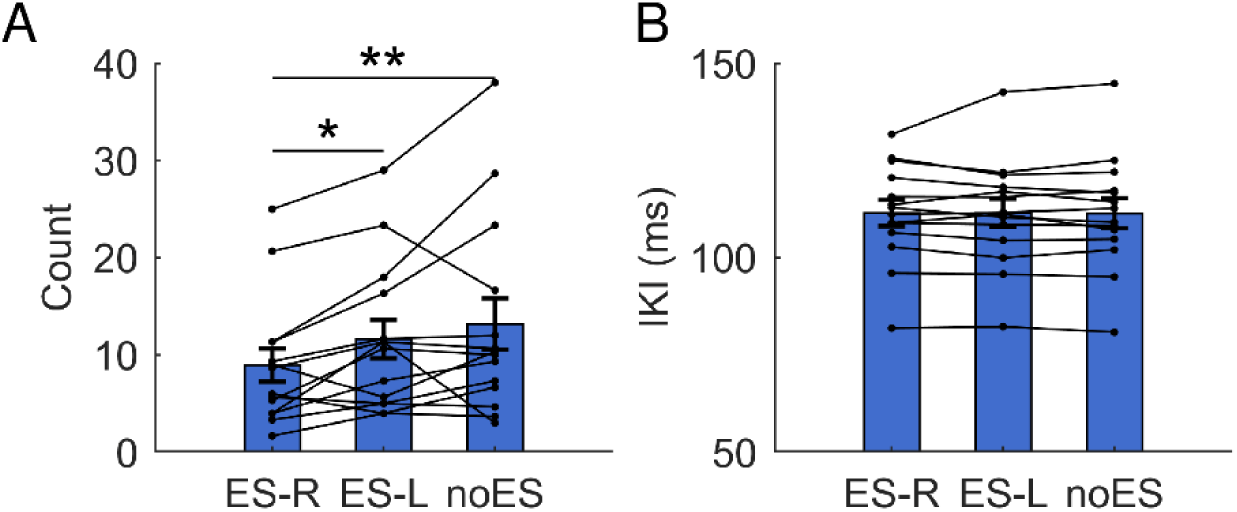
Effects of weak ulnar nerve stimulation on the performance of sequential finger movements. The group mean of the number of erroneous strikes due to incorrect fingering (A) and the interkeystroke interval (B) during the repetition of the finger sequential movement for 20 s. Mean±SEM. Small dots and thin lines indicate individual data. **, *: post hoc comparison, p<0.01, p<0.05.

## Discussion

The present study examined the experience-dependent plasticity of an inhibitory function in somatosensory processing by comparing expert pianists and nonmusicians. The series of five experiments demonstrated that pianists have superior ability to perceive the fast multifinger movements than nonmusicians and exhibit a unique somatosensory inhibition function in the S1 which can be activated by weak ulnar nerve stimulation. This weak stimulation activated surround inhibition circuits in the S1, increased the distance of cortical representation in the S1 across digits, and improved both the perception and execution of fast and complex multifinger sequential movements. These observations were absent in nonmusicians. Taken together, these findings demonstrated the experience-dependent plasticity in the inhibitory function of somatosensory processing in the S1 through extensive training of dexterous motor skills, which underlies experts’ ability to accurately perceive and control fast and complex movements.

### Inhibitory effect of weak ulnar nerve stimulation on somatosensory processes specifically in pianists

We examined the inhibitory function in the S1 using the paired-pulse paradigm of SEP. Previous studies demonstrated that repetitive TMS applied to the S1 and repetitive sensory stimulation modulate both tactile perception and PPS(*26*, *27*), and PPS was reduced by aging(*18*), indicating the ability of this inhibitory function to be plastically adapted by experiences and repetitive sensory inputs. A clear PPS was observed in the present study using the conventional paired-pulse method, in which the intensities of two stimuli were identical and close to the motor threshold in both the ulnar and median nerves. However, the group difference in the PPS was observed only when applying ulnar nerve stimulation with a CS intensity just below the PT. Previous studies demonstrated that weak tactile stimulation of the fingertips decreased BOLD activities in the S1 and increased the perceptual detection threshold for a subsequent tactile stimulus(*28*, *29*), suggesting selective activation of inhibitory interneurons in the S1. Animal studies also demonstrated a lower excitation threshold in inhibitory interneurons than in principal neurons in the S1(*30*, *31*). Although the present study is different from these previous studies in the site of stimulation, a weak stimulus would selectively activate inhibitory interneurons in the S1. In contrast, the conventional method may activate both excitatory and inhibitory neurons in the S1. Thus, we speculate that the excitatory activity evoked by intense stimulation may mask the group difference in the inhibitory effect.

In contrast, we did not find any group differences in the PPS in the median nerve at either stimulus intensity. The median and ulnar nerves at the wrist innervate different intrinsic hand muscles. The ulnar nerve at the wrist innervates muscles associated with all digits, while the median nerve at the wrist primarily innervates muscles associated with the thumb. The thumb is used frequently in daily life, is highly independent from the fingers anatomically and functionally(*32*, *33*), and has a unique cortical representation (*2*). Piano playing requires skilful control of the thumb and fingers(*12*). These requirements of all digits would induce experience-dependent neuroplasticity in pianists(*34*), which is a possible reason underlying the group difference in the PPS specific to the ulnar nerve in the present study.

### Weak ulnar nerve stimulation activates surround inhibition

An important role of somatosensory information processing is to decode which body part’s movement is encoded from the afferent information. Correction of ongoing movements based on incorrectly decoded sensory information can lead to the production of erroneous motor actions. This is a critical problem, particularly for experts, such as musicians and athletes, who need to control multiple body parts at high speed and with high accuracy. Previous studies demonstrated that cortical representation overlaps between multiple body parts in the sensorimotor cortex(*2*, *9*) and that some sensory inputs from different body parts converge into identical neurons(*3*, *4*). This may cause the sensory system to decode information from somatosensory inputs incorrectly when somatosensory information is inputted from different body parts within a short time period. To avoid this, somatosensory information must be processed independently for each body part. Here, we showed the superior ability to perceive the fast multifinger movements in pianists than nonmusicians. This suggests that the pianist’s nervous system processes somatosensory information from multiple fingers, interfering with each other to a smaller extent. We also found that pianists possess a unique inhibitory function that can modulate finger representation in somatosensory processing. The inhibitory function was activated by weak ulnar nerve stimulation and segregated the somatosensory processes between fingers. The latency of this effect was approximately 108-132 ms relative to the onset of passive finger movement. A previous study confirmed that the S1 is the origin of this component of SEP(*35*), indicating that the inhibitory activity evoked by weak ulnar nerve stimulation modulates neural activity in the S1. A putative mechanism behind this is the involvement of surround inhibition. We found that weak ulnar nerve stimulation enhanced surround inhibition as assessed by the dual stimulation technique. In addition, the stimulation lowered the perceptual threshold regarding the discrimination of the order of multifinger sequential movements. This perceptual enhancement and the amount of inhibition activated by the stimulation were correlated; pianists who had stronger inhibition showed larger perceptual enhancement. These effects were specific to the pianists, suggesting experience-dependent plasticity in this function. Taken together, pianists possess a unique surround inhibition function that sharpens the somatosensory processing of individual fingers, which prevents the interference of cortical activities across fingers and enables pianists to perceive fast multifinger movements correctly. One may wonder why the perceptual threshold did not differ between the two groups in Experiment 3. In this experiment, participants were asked to discriminate two movement sequences using the two-alternative forced choice method. In this procedure, participants need not process somatosensory information about all fingers, as they only need to perceive the last (or third) finger that moved. This simplicity of the task may reduce the requirement of segregated somatosensory processing of the individual fingers and thereby mask the difference in the threshold between groups. This idea is compatible with our finding of lower perceptual thresholds in pianists than non-musicians for the task used in Experiment 1, which required processing somatosensory information on all four fingers.

In addition to the perception, weak ulnar nerve stimulation enhanced the execution of fast and complex movements of multiple fingers. Here, we emphasize that the performance level of our participants was very high, as evidenced by only a few mistouches at a repetitive keystroke rate above 9 Hz. Surprisingly, such a highly trained motor skill was further enhanced by receiving weak ulnar nerve stimulation. The stimulation reduced the number of erroneous finger selections during movement without changing the movement speed. As discussed above, the weak ulnar nerve stimulation would activate the surround inhibition circuits in the S1. This would reduce interference of somatosensory information processing between the individual fingers during performing the task and thereby facilitate the execution of fast and complex movements. This indicates that the specialized inhibitory function in pianists plays a critical role in both accurate perception and execution of fast multifinger movements.

Taken together, the present study examined the experience-dependent plasticity of the inhibitory function of the somatosensory system. We first identified a pianist-specific somatosensory inhibitory function that can be activated by weak electrical stimulation of the ulnar nerve. This stimulation enhanced surround inhibition in the S1 and sharpened the representation of fingers in early somatosensory processing. Furthermore, both the perception and production of fast multifinger movements were improved by the stimulation. Taken together, these findings demonstrate the experience-dependent plasticity of surround inhibition function in the S1 and its functional role in the fast and complex movements of expert pianists.

## Acknowledgements

The present study was supported by a JSPS Grant-in-Aid for Transformative Research Areas B (20H04093) for M.H. and by JST Moonshot R&D (JPMJMS2012) for S.F.

## Materials and Methods

### Participants

In total, 71 healthy pianists and 57 healthy nonmusicians participated in this study. All of the pianists majored in piano performance at a musical conservatory and/or had extensive and continuous private piano training under the supervision of professional pianists and/or piano professors. Nonmusicians had no experience practising any musical instruments for over 5 years. All participants gave their written informed consent before engaging in the experiments. All experimental procedures were carried out in accordance with the Declaration of Helsinki and were approved by the ethics committees of the Sony Corporation.

### Experiment 1 Participants

Fifteen expert pianists and 15 nonmusicians participated in this experiment.

### Perception test

Participants wore an exoskeleton robot hand(*22*) on their right hand during the perception test. The robot hand can produce flexion and extension movements on the metacarpophalangeal joint of five digits. In this test, second to fifth digits were flexed passively with a randomized order except for the order of 2-3-4-5 and 5-4-3-2 (i.e., one of the 22 possible sequences was randomly chosen, 2: index finger, 3: middle finger, 4: ring finger, 5: little finger) at a constant angular velocity of 450 degree/s. The participants were asked to identify the order of passive finger movements and to respond to the perceived order by typing on a keyboard with their left hand using the same fingering as the perceived one. The interval between onsets of each finger flexion in the sequential passive movement varied across trials, which was determined by an adaptive Bayesian staircase method (ZEST)(*23*) to assess the perceptual threshold at which a participant was able to identify the order of a sequence of passive finger movements with 80% accuracy. Briefly, ZEST is an efficient method of measuring perceptual thresholds. We first specified a uniform distribution as the prior knowledge for the probability of threshold values and the Weibull function with a slope factor of 3.5 as the psychometric function. The probability density function (pdf) was updated every trial based on the result of the discrimination test, the stimulus intensity of that trial, and the psychometric function. The stimulus intensity and the final estimate of the threshold were chosen to correspond to the mean of the latest pdf. In this test, the chance level was 1/22 because the sequence at a given trial was randomly selected from 22 possible sequences. Therefore, we specified the gamma value in the Weibull function to this chance level. The test consisted of 50 trials.

### Experiment 2

#### Participants

Fourteen expert pianists and 14 nonmusicians participated in this experiment.

#### EEG recording and preprocessing

EEG signals were recorded from the CP3 electrode according to the international 10/20 system (actiCHamp plus and actiCAP snap, BrainProducts, Germany; impedance < 10 kΩ). The reference electrode was placed at Fz, and the sampling rate was 2.5 kHz. The offline analysis was performed using EEGLAB and a custom-made script using MATLAB. For offline analysis, EEG signals were preprocessed by bandpass filtering (0.5-300 Hz) and line noise removal (EEGLAB cleanLineNoise function).

#### Electrical stimulation

An electrical stimulus (a rectangular pulse of 1 ms duration) was applied to the median and ulnar nerves at the wrist using a constant current stimulator (DS4, Digitimer, Inc.) via a paired bar-type electrode. Before the experiment, we assessed the perceptual detection threshold (PT) of electrical stimulation for both nerves. The PT was defined as the lowest electrical current at which a participant reported sensation. Then, we determined the motor threshold (MT), the lowest electrical current that evokes a twitch of muscles innervated by each of the ulnar and median nerves (abductor pollicis brevis for the median nerve, first dorsal interosseous muscle for the ulnar nerve).

#### Paired-pulse suppression of somatosensory evoked potentials

The somatosensory evoked potentials (SEPs) were individually evoked by applying the electrical stimulus to each of the sensory nerves. To assess paired-pulse suppression (PPS), two electrical stimuli were applied to the nerve in succession. We set the intensity of the first stimulus, termed the conditioning stimulus (CS), at either just below the PT (PT-CS) or just below the MT (MT-CS). The second stimulus, termed the test stimulus (TS), was set at an intensity just below the MT to evoke a clear SEP. The interstimulus interval (ISI) between the CS and TS was set as 5, 10, 20, 30, 40, and 50 ms. In addition, there were conditions in which only PT-CS or only TS was presented. Therefore, in total, EEG responses were recorded for 14 stimulus conditions (6 ISI * 2 different intensities of CS, PT-CS-only, and TS-only) for each of the median and ulnar nerve conditions. Each stimulus condition was repeated 250 times. The different ISIs and intensities were presented stimulus-by-stimulus in a randomized order for each of the ulnar and median nerve stimulation conditions.

#### SEP analysis

The preprocessed EEG data were segmented in epochs −50 ms to 100 ms relative to the stimulus onset, and segments containing artefacts were removed (EEGLAB functions pop_jointprob.m, pop_rejkurt.m, both SD = 3, amplitude during pre-stimulus period > 50 μV). Then, segmented data were superimposed to increase the signal-to-noise ratio. For the paired-pulse SEP, potentials evoked by the CS may temporally overlap with potentials evoked by the subsequent TS. To remove this overlapping effect, we subtracted the SEP waveform obtained from the CS-only condition from that obtained from the paired-pulse conditions. Specifically, the SEP waveform in the PT-CS-only condition was subtracted from those obtained from the PT-CS conditions after adjusting the timing of stimulus onset, and the SEP waveform in the MT-CS conditions was subtracted from those obtained from the TS-only condition. We then determined that the negative peak component appeared approximately 20 ms after stimulus onset (N20) and that the positive peak appeared approximately 25 ms after stimulus onset (P25) and calculated the peak-to-peak amplitude between the N20 and P25 components.

### Experiment 3

#### Participants

Fourteen expert pianists and 14 nonmusicians participated in this experiment.

#### SEP recording

SEP was assessed using the same procedure and the same equipment as in Experiment 1.

#### Perception test

The participants wore an exoskeleton robot hand(*22*) on their right hand during the perception test. The robot hand can produce flexion and extension movements on the metacarpophalangeal joint of five digits. In this test, the finger joints of the robot hand were flexed passively with one of the predetermined sequences (1-3-2-5 or 1-3-5-2; 1: thumb, 2: index finger, 3: middle finger, 5: little finger) at a constant angular velocity of 450 degree/s. After the passive movements, the participants were asked to press one of the two keys put in front of them to identify the sequence in which their fingers were moved by the robot hand (i.e., two-alternative forced choice task). The interval between onsets of each finger flexion in the sequential passive movement varied across trials, which was determined by an adaptive Bayesian staircase method (ZEST)(*23*) to quantify the perceptual threshold at which a participant was able to discriminate the order of a sequence of passive finger movements with 80% accuracy. In this experiment, there were two conditions. In the first condition, a weak electrical pulse (intensity just below the PT, 1 ms square pulse) was applied to the ulnar nerve at the onset of each passive finger flexion (i.e., the stimulation was applied 4 times in a single passive sequential movement because the sequence used in this test consisted of 4 successive finger flexions). In the second condition, electrical stimulation was not provided. The two conditions appeared randomly in a trial-by-trial manner, and the corresponding pdf was updated based on the correctness of the answer. Each condition had 75 trials.

### Experiment 4

#### Participants

Fourteen pianists and 14 nonmusicians participated in this experiment.

#### EEG recording and preprocessing

EEG signals were recorded with 64 active electrodes according to the international 10/20 system (actiCHamp plus and actiCAP snap, BrainProducts, Germany; impedance < 10 kΩ). The reference electrode was placed at Fz, and the sampling rate was 1 kHz. The offline analysis was performed using EEGLAB and a custom-made script using MATLAB. For offline analysis, EEG signals were preprocessed by downsampling (250 Hz), low-cut filter (0.5 Hz), line noise removal (EEGLAB cleanLineNoise function), bad channel rejection, artefact subspace reconstruction (EEGLAB pop_clean_rawdata function, burst criterion: 40, window criterion: 0.25, burst rejection: on), channel interpolation, rereference to the average, and independent component analysis (ICA) with the Picard algorithm. Components extracted from the ICA (i.e., ICs) were classified using a machine learning algorithm implemented in the EEGLAB IC label function. ICs classified with less than 5% probability of brain activity and more than 50% probability as muscle activity were removed from the data.

#### Passive finger movements

One of the five metacarpophalangeal joints of the right hand was flexed from 0° to 45° at a constant speed of 450 degree/s using the exoskeleton robot hand. The same finger was extended from 45° to 0° at the same angular velocity 500 ms after flexion. In addition, there was another condition in which both the ring and little fingers were simultaneously flexed (i.e., dual condition). The interval of successive finger flexions was randomly determined from 2 s to 3 s. The number of passive movements was 200 for each digit and dual condition. In half of the trials, a weak electrical pulse (intensity just below the PT, 1 ms square pulse) was applied to the ulnar nerve at the wrist at the onset of each passive flexion (i.e., ES condition) using a constant current stimulator (DS4, Digitimer, Inc.) via a paired bar-type electrode. In contrast, no electrical stimulation was provided in the remaining trials (i.e., noES condition). To avoid the contamination of artefacts originating from eye movements in EEG data, we asked participants to close their eyes and rest during EEG recording.

#### Time-resolved representational similarity analysis

We used a time-resolved representational similarity analysis, where the pairwise dissimilarity distance of multichannel EEG signals between individual digits was calculated for each sample of the target time window. In this analysis, we used data obtained from the all sensors used in this study (i.e., 64 sensors). The preprocessed EEG signals were segmented by the onset of passive finger flexion (i.e., somatosensory stimulation) extended from −200 ms to 400 ms. We randomly divided the segments of each finger into 10 pseudo-blocks (i.e., each block contained a maximum of 10 segments per condition). Then, we estimated the SEP for each finger and condition in each pseudo-block using a linear model consisting of the finger (i.e., 5 fingers), condition (i.e., noES or ES), and their interaction as fixed effects. The model also includes the mean amplitude over the baseline period (i.e., −200-0 ms) and the interactions between the baseline and finger and between the baseline and condition as covariates(*36*). The estimated SEP amplitudes at each sample in each pseudo-block was prewhitened by the covariance of the residuals of the linear model (i.e., multivariate noise normalization) to minimize the effects of noise on the dissimilarity distance. The dissimilarity values of the prewhitened SEP amplitude patterns between all possible pairs of digits for each sample were measured using the cross-validated squared Euclidean distance (this is the same distance measure as the crossnobis distance reported in the previous study(*37*)). The dissimilarity of a sample t between digits i and j (i.e., 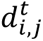) was obtained as follows:

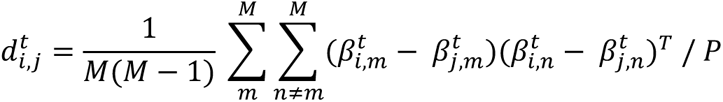

where M is the number of pseudo-blocks (10 in this experiment), 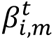 represents the prewhitened intersensor SEP amplitude pattern evoked by the i-th digit movement of sample t in the m-th pseudo-block, and P is the number of sensors (64 in this experiment). This analysis was repeated for every sample and digit pair. Furthermore, this procedure was repeated 20 times by randomly re-assigning the segments into 10 pseudo-blocks, and the resultant 20 dissimilarity time courses for each digit pair were averaged. The multivariate noise normalization and the cross-validated squared Euclidean distance were calculated according to previous studies(*37*, *38*). Due to the cross-validation procedure, the expected value of the distance was zero (or below) if two patterns were not significantly different from each other and larger than zero if the two patterns were different. Finally, we averaged the dissimilarity time courses across all digit pairs.

#### Cluster permutation test

To assess the interactive effect between the group and condition (i.e., ES and noES conditions) on the average dissimilarity time course, we performed the cluster permutation test. We first calculated the difference in the average dissimilarity time course between the ES and noES conditions for each group. Then, a two-sample t test was performed on the differential dissimilarities between the two groups for all samples. All samples with a p value larger than 0.05 were selected. Selected samples were clustered in connected sets according to temporal adjacency. Cluster-level statistics were calculated by taking the sum of the t values within every cluster. In the next step, we shuffled the group label for each participant. Then, a two-sample t test was performed on the differential dissimilarities between the shuffled groups for all samples. We identified the clusters and selected the cluster with the maximum cluster t value. This procedure was repeated 10,000 times, and we obtained a permutation distribution. Finally, we calculated the proportion of clusters that resulted in a larger t value in the permutation distribution than the observed ones. This proportion is the Monte Carlo significance probability, which also corresponds to a p value.

#### Somatosensory surround inhibition

We used the dual-SEP technique to assess somatosensory surround inhibition. The SEP waveform evoked by the simultaneous flexion of the ring and little fingers (i.e., dual SEP) was compared to the arithmetic sum of two SEPs (i.e., sum SEP) evoked by individual passive movement of the two fingers. We focused on the CP3 electrode, which was placed just above the sensorimotor area. The SEP evoked by passive movement is typically characterized by several components. A previous study demonstrated that the first component observed at approximately 100 ms originates from activities of the somatosensory cortex. Therefore, we quantified the peak amplitude of the first component and then compared it between the dual SEP and sum SEP.

### Experiment 5

#### Participants

Fourteen pianists participated in this experiment.

#### Sequential finger movement test

The pianists were asked to strike piano keys using their right hand in the following order: 1-3-4-2-3-5 (1: thumb, 2: index finger, 3: middle finger, 4: ring finger, 5: little finger). They repeated the sequence for 20 seconds as fast and accurately as possible during a trial. During this task, electrical pulses (1 ms square wave) with an intensity just below the PT were delivered to the right or left ulnar nerve at a frequency of 10 Hz. For the control condition, no stimulation was provided during the task. One stimulation device (DS4, Digitimer, Inc.) was used to stimulate the ulnar nerve on the right hand, and another device (DS7A, Digitimer, Inc.) was used to stimulate the ulnar nerve on the left hand. Both devices delivered electrical pulses via a pair of Ag/Cl electrodes placed on the wrist. Before the experiment, the pianists performed the task twice without any stimulation as a practice. Then, they performed the task 3 times for each condition in a randomized order (i.e., in a total of 9 trials). To avoid muscle fatigue, the intertrial interval was set at 2 min. We used a mute piano and instructed each participant to perform the task while listening to white noise from headphones worn on the ears and while closing their eyes.

### Statistical analysis

The data were analysed using a generalized linear mixed-effects (GLME) model implemented in the lme4 package(*39*) in R (https://www.r-project.org). Our model included the factors appropriate for the data with the group, stimulation condition, ISI, and nerve as fixed effects along with their interactions. Individual variability was assessed by including participants and participant*within factor interactions (when the number of within factors was 2 or more) as random effects. To determine the significance of the fixed effects, Type II Wald chi-square tests were performed using the ANOVA function in R with the car package(*40*). Multiple comparisons were performed based on the estimated marginal means calculated using the emmeans package(*41*) in R. The degrees of freedom of these comparisons were calculated using the Kenward-Roger method, while p values were adjusted using Hold’s method.

